# Does estimation methods affect on phosphorus equivalence value of phytase for layers and broiler chickens?

**DOI:** 10.1101/2020.09.26.314666

**Authors:** Azam Yousefi, Mojtaba Zaghari

## Abstract

Two experiments were performed for evaluating calibration curve (CC) and comparing negative and positive controls (CNP) as a major method for estimating of phytase phosphorus equivalence for layer and broiler chickens. In the first and second experiments, 360 Hy-line W-36 layer hens and 525 day-old Ross-308 broiler chickens were used in a complete randomized design, respectively. Evaluated methods were setting the two regression equations for NPP-supplemented and phytase supplemented treatments with two sub-methods, include calibration curve (CC) or exclude the amount of phosphorus content of basal diet (CC-BD) in calculation, and exploring enzyme equivalency by comparing phosphorus deficient diet as an negative and supplemented diet by inorganic phosphorus sources as a positive control group (CNP). Experiment one included nine treatments (200, 300, 400 and 500 FTU/kg phytase was added to a phosphorus deficient basal diet contained 0.12% AvP, the rest four treatments were included basal diet supplemented with 0.20, 0.27, 0.35 and 0.43% AvP). Experiment two included seven treatments (a basal P deficient diet contained 0.27% AvP, and two increasing levels of AvP, 0.32 and 0.37%, and four doses of phytase 200, 300, 400 and 500 FTU/kg added to basal diet). Each treatment in the both experiments replicated five times. Results indicated that methods of estimation had a significant effect on phosphorus equivalence estimation (P<0.0001). Fitted regression equations considering P content of basal diet (CC-BD) estimated rational values than those ignore it (CC) (0.161% *vs* 0.365% and 0.432% *vs* 0.564% for 500 FTU/kg phytase for broiler chicken and layer hens, respectively) (P<0.0001). On average, among three methods used, CC method had the highest estimated values both in broiler chickens and layer hens (p<0.0001). Regardless of mathematical method, there were different significant values for different strains (layer, 0.381% and broiler, 0.179%) (P<0.0001), but not for different traits served as response criteria (P>0.05). In conclusion, the phosphorus equivalent value of enzyme varies according to the estimation methods and strain. Hence, using matrix values of enzyme for accurate feed formulation depend on a variety of circumstances and decision making requires comprehensive information.

## DESCRIPTIONOF PROBLEM

Standard phytase activity defines as the amount of enzyme that releases 1 μmol of inorganic phosphate from a sodium phytate substrate per minute at pH 5.5 and 37 °C and expressed as FTU, FYT or OUT per Kg of feed. But, it can’t be an appropriate indicator to predict *in-vivo* efficiency of phytase. Because, many factors affect phytase functionality in practical nutrition (Bedford and Patridge, 2011). Dersjant-Li et al. (2019) reported that the pH optima range for various phyatses can be remarkably different. Phosphorus equivalency illustrates the potential of the enzyme to adding phosphorus to the dietor phosphorus contribution of a given unit of phytase *in-vivo*. Numerous studies have determined P equivalence of various phytases in poultry feeds. Interestingly, these values have been influenced by the source of phytase (Rodriguez et al., 1999 a,b; Tran et al., 2011), source of in-organic P (Li et al., 2015) P and Ca content of basal diet, Ca:P ratio of basal diet (Li et al., 2013), phytase inclusion rates in diets (Abd El-Hack et al., 2018),intended strain (Leskeand Coon, 1991) and finally, the manner of estimation (Dersjant-Li et al., 2019). The extent of phytase action is not limited to the P releasing. It is approved that supplementation of phytase in poultry diet, not only improves phosphorus availability but also the bioavailability of some other minerals, protein, amino acids and even energy (Jalal and Scheideler, 2001; Newkirk and Classen, 2001; Rutherfurd et al., 2004; Liu et al., 2009; Ghosh et al., 2016). Therefore, matrix value should estimates the releasing extent of the first limiting nutrient (i.e. P) and secondly Ca, Na, protein, AME and some other minerals in body using the recommended dose of enzyme.

Matrix values have been determined under controlled in-vivo experiments, however, the claimed nutrient saving values must be guaranteed by a significant degree of confidence. Overestimation or underestimation of equivalence values obtained for phytase may result in economic losses (Bedford et al., 2016).

Besides the variations resulted by different experimental assays adopted for P equivalence estimation (i.e. directly through digestibility tests or indirectly using a biological response criterion) (Bedford and Cowieson, 2020) or determinant factors (Ca and P content of basal diet, dietary fat content and …; Bedford et al., 2016), it seems that within a distinct manner of measurement, the method of P equivalence calculation is capable to overestimate or underestimate equivalence values.

In current study two performance trials fully described by Bedford and Cowieson (2020) had employed to determine the nutrient equivalence of a commercial phytase cocktail in both broiler chickens and layer hens. Three different methods within calibration curves as a major method, have adopted to calculate the P equivalence values of phytase in broiler chickens and layer hens.

## MATERIALS AND METHODS

### Experiment 1

Three hundred sixty 70-wk-old, W-36Hy-line layer hens were used in current experiment to estimate phosphorus equivalence of phytase. Layer hens were allotted to nine treatments and five replicates in a complete randomized design. A basal diet with 0.12% AvP was formulated and dietary treatments included four increasing levels of 0.07, 0.15, 0.21 and 0.31% NPP (equivalent to 0.20, 0.27, 0.35 and 0.43 % AvP, respectively) and four increasing doses of phytase 0.002, 0.003, 0.004 and 0.005 g/kg feed (equivalent to 200, 300, 400 and 500 FTU/kg). One unit FTU of phytase is defined as the quantity of enzyme, which liberates 1µmol of inorganic phosphate per minute from sodium phytate at pH 5.5 and 37 °C. The compositions of experimental diets are shown in table 1. Daily feed intake was 100 g/bird. All diets were iso-energetic and iso-nitrogenous by substituting an inert filler with DCP and phyase.

**Table 1.**
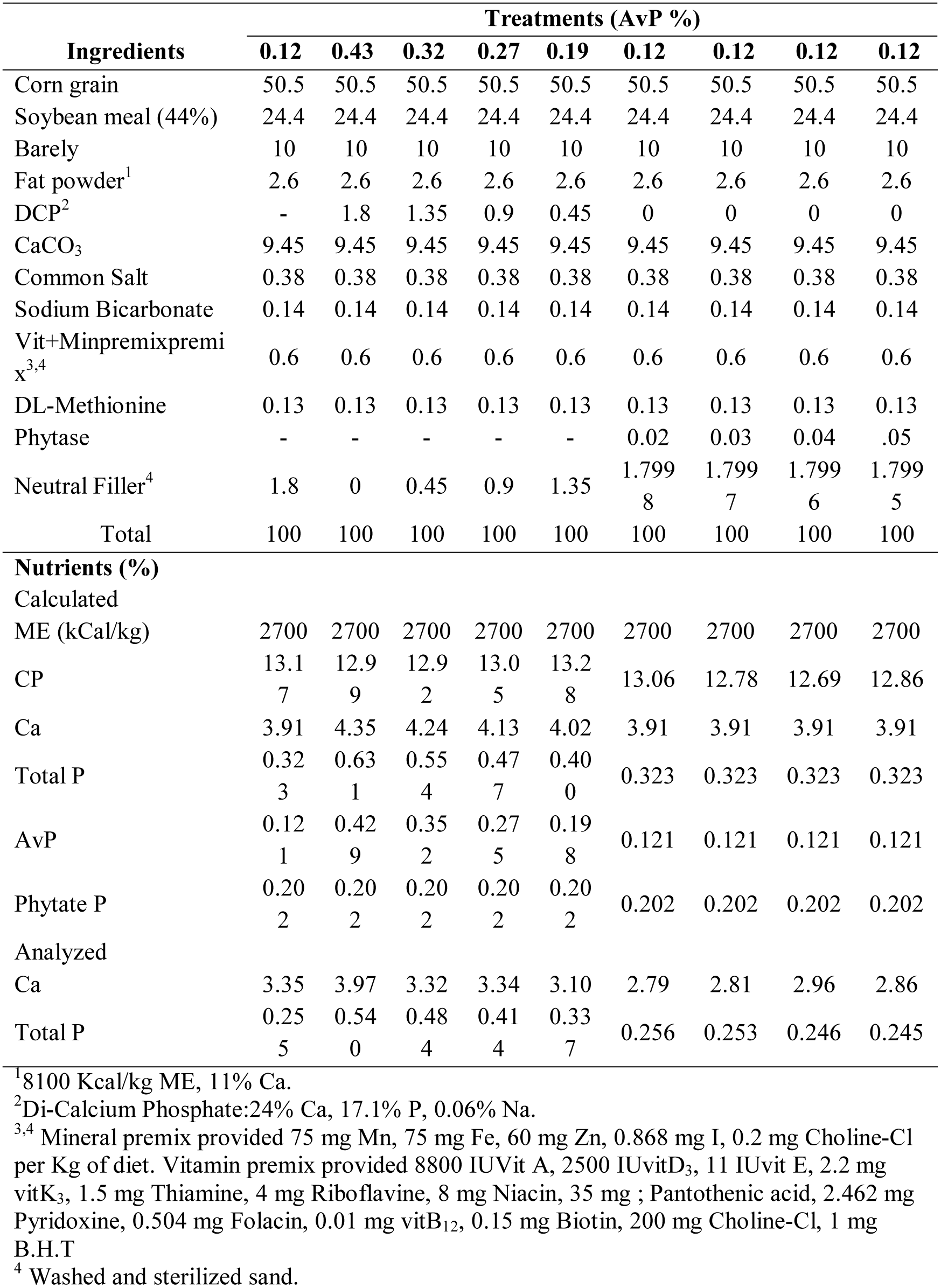
Diet composition and nutrient analysis of the experimental diet for layer hens.

During six weeks of experiment, total egg production and total saleable egg production were recorded daily and percentage hen-day egg production was calculated. All laid eggs were weighted once in a week. Two samples were selected and three different indicator locations on eggshell were measured, as stated by Zaghari (2009) and the mean value was reported as eggshell thickness. Egg mass was calculated as egg production rate×egg weight. Weekly feed intake (g) and egg mass (g) were used to calculate feed conversion ratio.

### Experiment 2

A total of 525 day-old male Ross-308 broiler chickens were allotted in seven treatments and five replicates of 15 birds in a complete randomized design. A basal diet was formulated to meet Ross 308 requirement except for phosphorus. Basal diet was supplemented by 1.1% mono-calcium phosphate to meet 56.25% of AvP requirement (i.e. 0.27%). Dietary treatments 1 and 2 were supplemented with 1.1 and 0.85% mono-calcium phosphate to provide 0.37 and 0.32 % AvP respectively. Diets 4 through 7 contained different levels of phytase (200, 300, 400 and 500 FTU/kg). Table 2 represents the ingredients and diet compositions. Broiler chickens were fed with a single diet from 1 to 28 days of age. Weight gain and feed intake were measured on days 7, 14, 21 and 28.Feed conversion ratio was calculated for weekly recorded data.

**Table 2.**
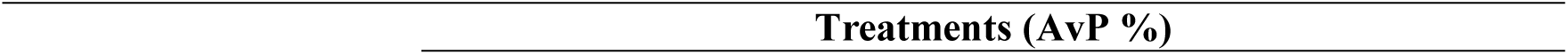
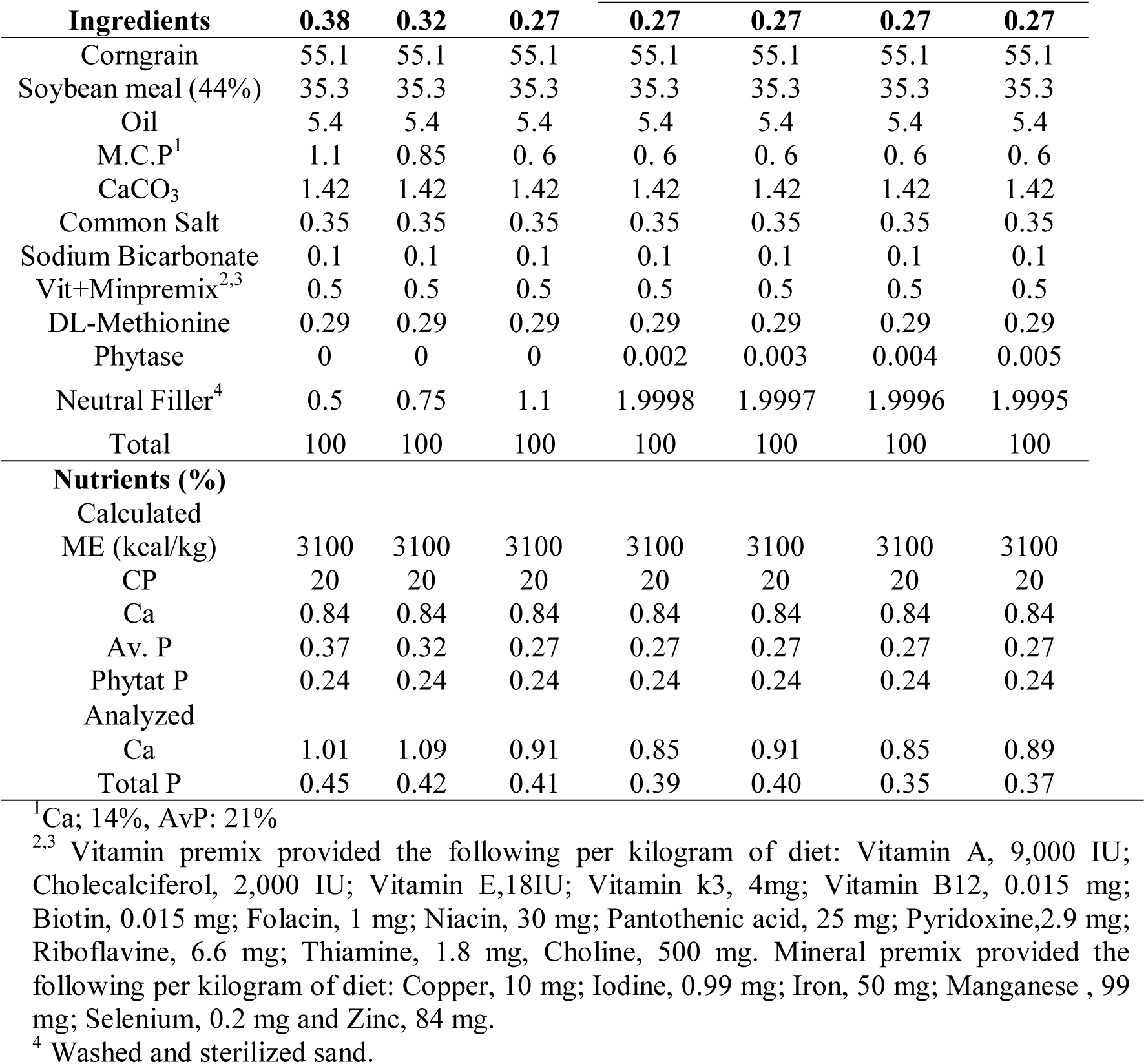
Diet composition and nutrient analysis of the experimental diet for broiler chickens.

| **Ingredients** | **0.38** | **0.32** | **0.27** | **0.27** | **0.27** | **0.27** | **0.27** |
| --- | --- | --- | --- | --- | --- | --- | --- |
| Corngrain | 55.1 | 55.1 | 55.1 | 55.1 | 55.1 | 55.1 | 55.1 |
| Soybean meal (44%) | 35.3 | 35.3 | 35.3 | 35.3 | 35.3 | 35.3 | 35.3 |
| Oil | 5.4 | 5.4 | 5.4 | 5.4 | 5.4 | 5.4 | 5.4 |
| M.C.P<sup>1</sup> | 1.1 | 0.85 | 0.6 | 0.6 | 0.6 | 0.6 | 0.6 |
| CaCO<sub>3</sub> | 1.42 | 1.42 | 1.42 | 1.42 | 1.42 | 1.42 | 1.42 |
| Common Salt | 0.35 | 0.35 | 0.35 | 0.35 | 0.35 | 0.35 | 0.35 |
| Sodium Bicarbonate | 0.1 | 0.1 | 0.1 | 0.1 | 0.1 | 0.1 | 0.1 |
| Vit+Minpremix<sup>2,3</sup> | 0.5 | 0.5 | 0.5 | 0.5 | 0.5 | 0.5 | 0.5 |
| DL-Methionine | 0.29 | 0.29 | 0.29 | 0.29 | 0.29 | 0.29 | 0.29 |
| Phytase | 0 | 0 | 0 | 0.002 | 0.003 | 0.004 | 0.005 |
| Neutral Filler<sup>4</sup> | 0.5 | 0.75 | 1.1 | 1.9998 | 1.9997 | 1.9996 | 1.9995 |
| Total | 100 | 100 | 100 | 100 | 100 | 100 | 100 |
| **Nutrients (%)** |  |  |  |  |  |  |  |
| Calculated |  |  |  |  |  |  |  |
| ME (kcal/kg) | 3100 | 3100 | 3100 | 3100 | 3100 | 3100 | 3100 |
| CP | 20 | 20 | 20 | 20 | 20 | 20 | 20 |
| Ca | 0.84 | 0.84 | 0.84 | 0.84 | 0.84 | 0.84 | 0.84 |
| Av. P | 0.37 | 0.32 | 0.27 | 0.27 | 0.27 | 0.27 | 0.27 |
| Phytat P | 0.24 | 0.24 | 0.24 | 0.24 | 0.24 | 0.24 | 0.24 |
| Analyzed |  |  |  |  |  |  |  |
| Ca | 1.01 | 1.09 | 0.91 | 0.85 | 0.91 | 0.85 | 0.89 |
| Total P | 0.45 | 0.42 | 0.41 | 0.39 | 0.40 | 0.35 | 0.37 |

<sup>1</sup>Ca; 14%, AvP: 21%
<sup>2,3</sup> Vitamin premix provided the following per kilogram of diet: Vitamin A, 9,000 IU; Cholecalciferol, 2,000 IU; Vitamin E, 18 IU; Vitamin K<sub>3</sub>, 4 mg; Vitamin B<sub>12</sub>, 0.015 mg; Biotin, 0.015 mg; Folic acid, 1 mg; Niacin, 30 mg; Pantothenic acid, 25 mg; Pyridoxine, 2.9 mg; Riboflavin, 6.6 mg; Thiamine, 1.8 mg; Choline, 500 mg. Mineral premix provided the following per kilogram of diet: Copper, 10 mg; Iodine, 0.99 mg; Iron, 50 mg; Manganese, 99 mg; Selenium, 0.2 mg and Zinc, 84 mg.
<sup>4</sup> Washed and sterilized sand.

### Statistical Analysis

The GLM procedure and Duncan multiple range test of SAS software (2004) were adopted to analyze data means. Statistical significance was determined at (P<0.05).Two regression equations (calibration curves) were created for two classes of treatments (NPP-supplemented treatments and phytase-supplemented treatments) for both laying hens and broiler chickens. Three different methods were used to calculate phosphorus release values.

Method one (Calibration Curve (CC)): Phosphorus equivalence was calculated by putting Y=treatment mean values into regression equations created for NPP-supplemented treatments as described by Fernandez et al., (2019) and solved as follow:

Linear function: Y=a+bX

_BW 28_= 720.093 + 1485.72 Phosphorus

Y_BW 28_= 1357.55 (Treatment mean, supplemented with 500 FTU/kg phytase)

Phosphorus=(1357.55-720.093)/1485.72

Phosphorus=0.429

Method two (Calibration Curve-Basal Diet Phosphorus (CC-BD)): Phosphorus equivalency was calculated by setting the two regression equations equal according to following procedure as described by Zaghari et al., (2008):

Y_BW 28_= 720.093 + 1485.72 Phosphorus,

Y_BW 28_= 1129.80 + 47000 Enzyme,

720.093 + 1485.72 Phosphorus=1129.80 + 47000 Enzyme

1485.72 Phosphorus=409.71 + 47000 Enzyme

Phosphorus=0.275 + 31.63 Enzyme

Phosphorus=0.275 + 31.63 (0.005)

Phosphorus=0.433 – (Phosphorus content of basal diet)

Phosphorus=0.433 – (0.27)

Phosphorus=0.164

Method three (Positive Control-Negative Control (CNP)): The third method is the product of the difference in AvP content between negative and positive controls, multiplied by the percentage of performance improvement of the phytase supplemented treatment compared to the positive control.

## RESULTS AND DISCUSSION

### Experiment 1

Effects of different levels of AvP and phytase on layer hens performance and egg quality are shown in table 3 Dietary treatments had no significant effects on egg production and egg quality variables during weeks 70 to 73 (P>0.05). Different levels of AvP and phytase led to improvements in FCR at weeks 74 and 75 (P<0.05). At week 75,number of produced egg, egg production percentage, saleable egg percentage and FCR in treatments with graded levels of AvP, were significantly different from negative control (without NPP and phytase) (P<0.05). Layer hens fed with 500 FTU/kg diet exhibited egg production percentage (EPP), saleable egg production percentage (SEP) and FCR equal to the birds fed positive control (0.43% AvP) (P< 0.05), as expressed previously by Shet et al., (2017). But, such an effect was not seen in treatments fed lower doses of phytase. The above results agree with Um and Paik, (1999) and Shet et al., (2017) who have reported in very low phosphorus diets (approximately 0.12 % AvP), higher dose of phytase (500 FTU/kg) could maintain laying performance without supplemental NPP, while lower doses (e.g 250 FTU/kg) were capable to perform moderate improvement of performance in diets that met 50% of AvP requirements.

**Table 3.** Effects of different levels of AvP and phytase on laying hen performance and egg quality (week 75)

| Variables | Treatments |  |  |  |  |  |  |  |  |  | SEM | P-value |
| --- | --- | --- | --- | --- | --- | --- | --- | --- | --- | --- | --- | --- |
|  | AvP<br>Phytase (FTU/kg) | 0.12 | 0.43 | 0.32 | 0.27 | 0.19 | 0.12 | 0.12 | 0.12 | 0.12 |  |  |
|  |  | - | - | - | - | - | 200 | 300 | 400 | 500 |  |  |
| **Total Egg** |  | 20.29<sup>c</sup> | 41.20<sup>a</sup> | 36.80<sup>ab</sup> | 37.20<sup>ab</sup> | 32.20<sup>bc</sup> | 34.80<sup>b</sup> | 37.40<sup>ab</sup> | 35.20<sup>b</sup> | 41.20<sup>a</sup> | 1.80 | 0.0006 |
| **Egg Production (%)** |  | 52.14<sup>c</sup> | 77.65<sup>a</sup> | 65.71<sup>b</sup> | 66.42<sup>b</sup> | 63.49<sup>b</sup> | 62.14<sup>b</sup> | 66.78<sup>b</sup> | 64.74<sup>b</sup> | 77.14<sup>a</sup> | 3.385 | 0.0002 |
| **SEP<sup>2</sup>** |  | 47.50<sup>d</sup> | 74.13<sup>ab</sup> | 64.64<sup>bc</sup> | 63.57<sup>bc</sup> | 60.07<sup>c</sup> | 57.85<sup>c</sup> | 64.28<sup>bc</sup> | 62.24<sup>c</sup> | 75.61<sup>a</sup> | 3.581 | 0.0001 |
| **FCR** |  | 3.32<sup>a</sup> | 2.17<sup>cd</sup> | 2.59<sup>bc</sup> | 2.56<sup>bc</sup> | 2.80<sup>b</sup> | 2.63<sup>b</sup> | 2.54<sup>bc</sup> | 2.56<sup>bc</sup> | 2.13<sup>d</sup> | 0.134 | <0.0001 |
| **Egg weight (g)** |  | 58.27 | 59.63 | 59.94 | 59.72 | 57.14 | 61.17 | 59.03 | 60.88 | 61.39 | 1.008 | 0.132 |
| **Yolk (%)<sup>3</sup>** |  | 28.27 | 29.73 | 28.14 | 28.52 | 28.09 | 27.76 | 28.47 | 28.80 | 28.32 | 0.0696 | 0.732 |
| **Egg shell thickness (mm)** |  | 0.344<sup>c</sup> | 0.366<sup>ab</sup> | 0.362<sup>abc</sup> | 0.362<sup>abc</sup> | 0.353<sup>bc</sup> | 0.372<sup>a</sup> | 0.352<sup>bc</sup> | 0.358<sup>abc</sup> | 0.366<sup>ab</sup> | 0.0057 | 0.045 |

<sup>a,b,c,d</sup> Means within a row with different superscripts differ (P<0.05).
<sup>1</sup>Egg Production Percentage
<sup>2</sup> (Yolk weight/egg weight)\* 100

On the other hand, the insignificant effects of phytase in P deficient diets during weeks 70 to 73 might be due to the releasing of Ca and P from medullary bone into blood stream (Whitehead and Fleming, 2000), which have decreased the efficacy of dephytinization. Fernandez et al., (2019) have stated at time lag the medullary bone resources compensate P and Ca requirements for egg production.

Table 4 shows the phosphorus equivalences of phytase in layer hens at week 75 got from three different methods i.e. solving of the regression equations with or without considering P content of basal diet and by comparing phosphorus contents of positive and negative controls. Phosphorus equivalences in the second method were calculated by subtracting the amount of available phosphorus of basal diet (i.e. 0.12%) from obtained values. Eggshell thickness, FCR, total egg production, egg production percentage and total saleable egg percentage showed a greater relationship with their respective regression equations compared to egg quality variables. The most dependent variable to P and phytase levels was FCR, (R^2^=0.53 and 0.67).

**Table 4.** Regression equations and estimated nutrient equivalency values of phytasein layer hens at week 75.

|  | **Egg shell**<br>**Thickness** | **FCR** | **Saleable egg percentage** | **Egg Production Percentage** |  |  |
| --- | --- | --- | --- | --- | --- | --- |
| **Equation** | Y=0.29+0.13P | Y = 3.99 - 3.27 P | Y=32.085+75.50P | Y=37.53+69.57P |  |  |
| **R<sup>2</sup>** | 0.40 | 0.53 | 0.52 | 0.45 |  |  |
| **P** | 0.0007 | <0.0001 | <0.0001 | 0.0002 |  |  |
| **Equation** | Y=0.33+7.45E | Y= 3.23 - 211.70 E | Y=47.38+5041.21E | Y=52.28+4395.27E |  |  |
| **R<sup>2</sup>** | 0.36 | 0.67 | 0.58 | 0.62 |  |  |
| **P** | 0.0015 | <0.0001 | <0.0001 | <0.0001 |  |  |
| **Phytase (FTU/kg)** | **P equivalence (Method CC)<sup>1</sup>** |  |  |  |  |  |
| **200** | 0.500 | 0.382 | 0.341 | 0.353 |  |  |
| **300** | 0.500 | 0.452 | 0.426 | 0.420 |  |  |
| **400** | 0.500 | 0.393 | 0.437 | 0.434 |  |  |
| **500** | 0.568 | 0.553 | 0.568 | 0.569 |  |  |
| **Phytase (FTU/kg)** | **P equivalence (Method CC-BD)<sup>2</sup>** |  |  |  |  |  |
| **200** | 0.298 | 0.238 | 0.170 | 0.215 |  |  |
| **300** | 0.356 | 0.303 | 0.240 | 0.187 |  |  |
| **400** | 0.413 | 0.367 | 0.306 | 0.341 |  |  |
| **500** | 0.470 | 0.422 | 0.372 | 0.404 |  |  |
| **Phytase (FTU/kg)** | **P equivalence (Method CNP)<sup>3</sup>** |  |  |  |  |  |
| **500<sup>4</sup>** | 0.293 | 0.313 | 0.314 | 0.308 | **P-Value** | **SEM** |
| **Average P**<br>**equivalence<sup>5</sup>** | 0.433 | 0.359 | 0.352 | 0.480 | 0.3802 | 0.0354 |

<sup>1</sup> Calibration Curve: Calculated by solving obtained regression equations for P by Y=treatment means.
<sup>2</sup> Calibration Curve-Basal Diet phosphorus: Calculated by setting the regression equations for P equal with those created for phytase, followed by subtracting phosphorus content of basal diet.
<sup>3</sup> Positive Control-Negative Control: Difference in Av. P content between negative and positive controls multiplied by the percentage of performance improvement of the phytase supplemented treatment compared to the positive control.
<sup>4</sup> Only 500 FTU/kg phytase resulted in identical performance to the positive control (0.43% AvP) for all response criteria.
<sup>5</sup> Average for all response criteria.

The amount of released P g^-1^ of phytase in 200 FTU/kg supplemented treatment was lower than 500 FTU/kg. These findings are in accordance with Fernandez et al. (2019) and Vieira et al. (2015) who have reported that phytase P release values increased with increasing the dosage of phytase.

Using eggshell thickness as the response parameter resulted in higher P equivalent values compared to other response criteria in methods CC and CC-BD but not in CNP. Estimated P equivalences got from three different methods is a little higher than the values of some other studies performed on laying hens (Simons and Versteegh, 1992; 1993; Waldroup, 1999), which could be attributed to the difference in the method of determination and experimental assays (digestibility trails *vs* performance trails) (Dersjant-Li et al., 2019) or different adopted response criteria (Adedokun et al., 2004), diet ingredients (Francesch et al., 2005), phytase type (Igbasan et al., 2000; Selle and Ravindran, 2007; Ribeiro et al., 2016), phosphorus source (Li et al., 2015), age of examination (Bedford and Cowieson, 2020) and protein and energy effect of phytase (Ravindran et al., 1999; 2000;Nahm, 2002; Liu et al., 2009). The later item needs more attention when interpreting the P equivalence of phytase. Because, phytase activity is not limited to the liberating phosphors, but it may influence performance by the ways independent to phytate-bound P release (Wu et al., 2004), therefore it probably results in over-estimating of P equivalence of a given phytase.

In the case of current study, it sounds that supplementing of a P deficient barley-based layer hen diet with phytase, resulted in higher mean P absorption as stated by Francesch et al., (2005) than those studies consisted of maize. More over, there are evidences of a complementary effect between intrinsic phytase of barley and supplemental phytase (Zyla, 1993; Näsi et al., 1999).

### Experiment 2

Table 5 represents the effects of graded levels of NPP and phytase on performance variables in broiler chickens. Growth performance showed no significant differences between dietary treatments at 7 d of age (P>0.05). Body weight gain and feed intake significantly influenced by dietary treatments during 14 to 28 d of age (P<0.05). Phytase supplementation (200 to 500 FTU/kg) recovers weight gain to the 0.21 g/kg NPP level at 14 and 21 d of age. But at 28 d of age, only 300-500 FTU/kg of phytase showed insignificant differences compared to 0.21 g/kg NPP supplemented (0.37% AvP) treatment. Feed conversion ratio didn’t show any significant difference at 14 and 28 d of age.

**Table 5.** Effects of different levels of AvP and phytase on broiler chickens growth performance (7-21 d of age)

| AvP (%)<br>Phytase (FTU/kg)<br>Variables | Treatments |  |  |  |  |  |  | SEM | P-value |
| --- | --- | --- | --- | --- | --- | --- | --- | --- | --- |
|  | 0.27<br>- | 0.32<br>- | 0.38<br>- | 0.27<br>200 | 0.27<br>300 | 0.27<br>400 | 0.27<br>500 |  |  |
|  |  |  | 0.38 |  |  |  |  |  |  |
| Weight Gain7-d (g) | 93.33 | 94.13 | 97.46 | 96.53<sup>a</sup> | 103.46 | 90.80 | 99.23 | 2.705 | 0.076 |
| Feed Intake 7-d (g) | 113.86 | 114.66 | 115.73 | 115.73 | 121.46 | 114.40 | 119.22 | 2.375 | 0.262 |
| FCR 7-d | 1.22 | 1.22 | 1.19 | 1.20 | 1.17 | 1.26 | 1.20 | 0.018 | 0.053 |
| Weight gain 14-d (g) | 291.07<sup>c</sup> | 314.80<sup>b</sup> | 317.47<sup>b</sup> | 317.20<sup>b</sup> | 341.33<sup>a</sup> | 319.60<sup>ab</sup> | 335.73<sup>ab</sup> | 7.340 | 0.0019 |
| Feed Intake 14-d (g) | 385.60<sup>c</sup> | 404.00<sup>bc</sup> | 423.73<sup>ab</sup> | 418.00<sup>ab</sup> | 425.60<sup>ab</sup> | 419.47<sup>ab</sup> | 432.80<sup>a</sup> | 7.690 | 0.0038 |
| FCR 14-d | 1.32 | 1.28 | 1.33 | 1.32 | 1.28 | 1.24 | 1.31 | 0.0205 | 0.1048 |
| Weight gain 21-d (g) | 613.11<sup>d</sup> | 654.13<sup>c</sup> | 702.80<sup>ab</sup> | 673.18<sup>bc</sup> | 720.53<sup>a</sup> | 702.44<sup>ab</sup> | 743.72<sup>a</sup> | 13.512 | <0.0001 |
| Feed Intake 21-d (g) | 853.91<sup>d</sup> | 893.20<sup>cd</sup> | 972.80<sup>a</sup> | 913.83<sup>bc</sup> | 958.27<sup>ab</sup> | 961.08<sup>ab</sup> | 996.67<sup>a</sup> | 18.411 | <0.0001 |
| FCR 21-d | 1.39<sup>a</sup> | 1.36<sup>abcd</sup> | 1.32<sup>ab</sup> | 1.35<sup>bcd</sup> | 1.33<sup>d</sup> | 1.37<sup>abc</sup> | 1.34<sup>cd</sup> | 0.0103 | 0.0025 |
| Weight gain 28-d (g) | 1090.83<sup>b</sup> | 1142.35<sup>b</sup> | 1225.07<sup>a</sup> | 1148.93<sup>b</sup> | 1263.47<sup>a</sup> | 1274.57<sup>a</sup> | 1292.27<sup>a</sup> | 25.031 | <0.0001 |
| Feed Intake28-d (g) | 113.86 | 114.66 | 115.73 | 115.73 | 121.46 | 114.40 | 119.22 | 33.559 | 0.262 |
| FCR 28-d | 1.43 | 1.45 | 1.44 | 1.43 | 1.39 | 1.4. | 1.42 | 0.0288 | 0.743 |

<sup>a,b,c,d</sup> Means within a row with different superscripts differ (P<0.05).

Table 6 shows the phosphorus equivalences (%) of different levels of phytase in broiler chickens using linear regression equations. Phosphorus equivalence values for broiler chickens follow the same principles as layer hens, (i.e. CC-BD, CC and CNP methods). Regression of body weight gain (BWG) on dependent variables at 14, 21 and 28 d of age, had higher R^2^ values than other variables, therefore the P equivalences were calculated for BWG as a response criteria. Potter (1988) introduces body weight gain and toe ash percentage as the best indicators for P equivalency measurement. Data showed that regardless of the method of calculating, the average P equivalence for BWG at different ages, were not significantly different (P>0.05) and ranged from 0.211 to 0.218% of AvP. However, when values compared individually based on the method of calculation, data were still in the range of previous studies, who showed that the amount of available P released by phytase ranged from 0.24 to 0.26 % (Plumstead et al., 2013), 0.035 to 0.208 % (Han et al., 2009) and 0.07 to 0.12% (Jendza et al., 2006) in broiler diets. Differences in type of diet, P content of basal diet, phytase type and the manner of experimental assay (digestibility trails or performance trails) might be responsible for the differences in calculated values between various studies. However, the results of current study (method CC-BD) were not entirely out of the range of the values by other performance experiments carried out broiler chickens.

**Table 6.** Regression equations and estimated phosphorus equivalence values of phytase in broiler chickens (14-21 d of age)

|  | Variables |  |  |  |  |
| --- | --- | --- | --- | --- | --- |
|  | BWG14 d | BWG21 d | BWG28 d |  |  |
| **Equation** | Y=270.74+39.44P | Y=419.49+888.86P | Y=720.09+1485.72P |  |  |
| **R<sup>2</sup>** | 0.378 | 0.713 | 0.695 |  |  |
| **P** | 0.0147 | <0.0001 | <0.0001 |  |  |
| **Equation** | Y=354.122+8253.24E | Y=668.82+24786E | Y=1129+47000E |  |  |
| **R<sup>2</sup>** | 0.342 | 0.60 | 0.762 |  |  |
| **P-value** | 0.0021 | <0.0001 | <0.0001 |  |  |
| **Enzyme levels (FTU/kg)** | **P equivalence (Method CC)<sup>1</sup>** |  |  |  |  |
| **200** | 0.177 | 0.285 | 0.289 |  |  |
| **300** | 0.269 | 0.339 | 0.366 |  |  |
| **400** | 0.186 | 0.318 | 0.373 |  |  |
| **500** | 0.247 | 0.365 | 0.385 |  |  |
| **Enzyme levels (FTU/kg)** | **P equivalence (Method CC-BD)<sup>2</sup>** |  |  |  |  |
| **200** | 0.076 | 0.066 | 0.070 |  |  |
| **300** | 0.107 | 0.094 | 0.100 |  |  |
| **400** | 0.139 | 0.122 | 0.132 |  |  |
| **500** | 0.170 | 0.150 | 0.164 |  |  |
| **Enzyme levels (FTU/kg)** | **P equivalence (Method CNP)<sup>3</sup>** |  |  |  |  |
| **300** | 0.108 | 0.102 | 0.103 |  |  |
| **400** | 0.101 | 0.100 | 0.104 |  |  |
| **500** | 0.106 | 0.106 | 0.105 | **P-value** | **SEM** |
| **Average P equivalence** | 0.136 | 0.185 | 0.199 | 0.977 | 0.0434 |

<sup>1</sup>Calibration Curve: Calculated by solving obtained regression equations for P by Y=treatment means.
<sup>2</sup> Calibration Curve-Basal Diet phosphorus: Calculated by setting the regression equations for P equal with those created for phytase, followed by subtracting phosphorus content of basal diet. <sup>3</sup>Positive Control-Negative Control: Difference in Av. P content between negative and positive controls multiplied by the percentage of performance improvement of the phytase supplemented treatment compared to the positive control.
Treatments supplemented with 300-500 FTU/kg phytase resulted in identical performance to the positive control (0.37% AvP) for all response criteria. <sup>5</sup>Average for all response criteria.

Table 7 represents the comparison of three different methods adopted for calculating phosphorus equivalences of phytase and average values obtained for different strains. For each phytase level, methods of calculating gave P equivalences, which were significantly different (P<0.0001). In broiler chickens, average phosphorus equivalence of 500 FTU/kg of phytase for BWG response (14, 21 and 28 days old), ranged from around 0.105 for CNP to 0.385 for CC methods. While, the phosphorus equivalences of 400 FTU/kg and 300 FTU/kg for BWG were in the range of 0.101 and 104 for CNP to 0.373and 0.366 for CC, respectively. In layer hens, the comparison of P equivalences values obtained by three methods of calculation, carried out only at the level of 500 FTU/kg phtytase. The CNP provided the lowest P equivalences in all phytase doses (P<0.0001). Comparison of P equivalences obtained by different methods was illustrative of underestimation of values obtained by CNP method in both layer hens and broilers. On the other hand, values obtained by CC method (without subtracting P content of basal diet) might be conflicting and may overestimates the P equivalence of phytase, because theoretically, it exceeded phytate phosphorus content of basal diet (i.e. 0.202% in layers and 0.24% in broilers). It may be concluded that CC method estimates total P release value in phytase-supplemented treatments, while it hasn’t subtract the P content of basal diet. Moreover, both in CC and CNP methods, the statistical influences of other doses of phytase have been ignored, when P equivalence of a given dose is calculated. Therefore, it is not surprising that calculated values are not supported by phytate content of basal diet. Bedford and Cowieson (2020) have stated that calculating of P equivalence of phytase through CNP method, may not be as accurate as using multiple calibration curves, because it strictly depends on difference of P content of negative control and positive control and real P requirements. Another interesting result was the higher P equivalences values obtained for layers compared with broilers. Average P release values of 500 FTU/kg phyatse were 0.433 and 0.230% in broilers and layers, respectively. This might be due to the nature of basal diet in two different experiments. Available phosphorus content of layer hens basal diet was approximately 3.5 times lower than recommended P requirements at this age (0.121% *vs* 0.40 to 0.42%). Therefore, the slopes for egg production equation derived from NPP-supplemented treatments in current study, were slightly higher than slopes derived for data reported by Fernandez et al., (2019) (69.75 *vs* 67.6). Consequently, the slopes for egg production equation created for phytase-supplemented treatments will increase exponentially and results seems that CC-BD method in the greater values, when equations set equal to obtain P equivalence of phytase. Therefore, obtained values may not be representative of commercial status performed by the end user.

**Table 7.**
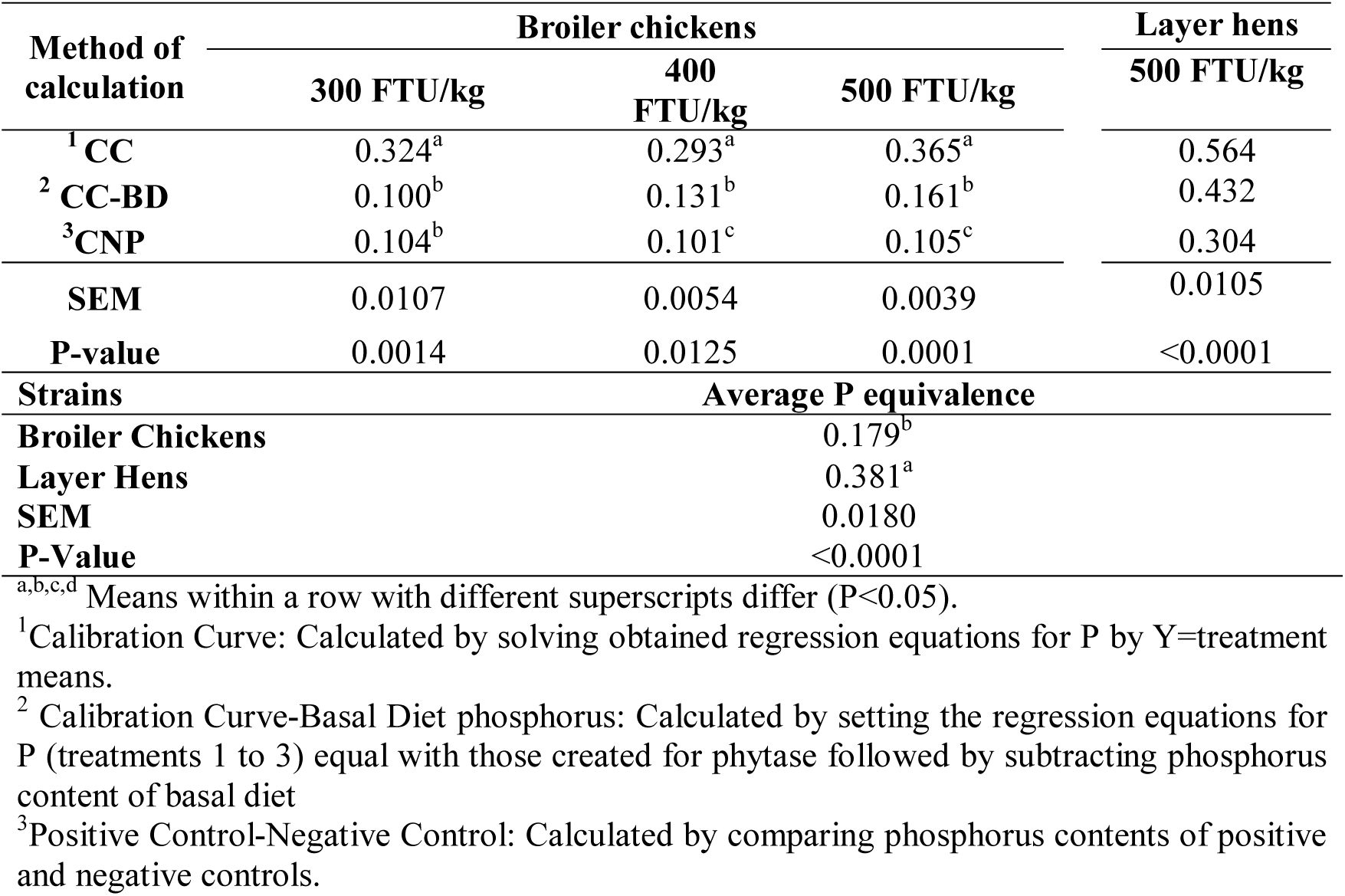
Comparison of different methods for estimating of phosphorus equivalence (contribution in the diet) values of phytase in different strains.

| Method of calculation | Broiler chickens |  |  | Layer hens |
| --- | --- | --- | --- | --- |
|  | 300 FTU/kg | 400 FTU/kg | 500 FTU/kg | 500 FTU/kg |
| <sup>1</sup>CC | 0.324<sup>a</sup> | 0.293<sup>a</sup> | 0.365<sup>a</sup> | 0.564 |
| <sup>2</sup>CC-BD | 0.100<sup>b</sup> | 0.131<sup>b</sup> | 0.161<sup>b</sup> | 0.432 |
| <sup>3</sup>CNP | 0.104<sup>b</sup> | 0.101<sup>c</sup> | 0.105<sup>c</sup> | 0.304 |
| SEM | 0.0107 | 0.0054 | 0.0039 | 0.0105 |
| P-value | 0.0014 | 0.0125 | 0.0001 | <0.0001 |
| **Strains** | **Average P equivalence** |  |  |  |
| **Broiler Chickens** | 0.179<sup>b</sup> |  |  |  |
| **Layer Hens** | 0.381<sup>a</sup> |  |  |  |
| **SEM** | 0.0180 |  |  |  |
| **P-Value** | <0.0001 |  |  |  |

<sup>a,b,c,d</sup> Means within a row with different superscripts differ (P<0.05).
<sup>1</sup>Calibration Curve: Calculated by solving obtained regression equations for P by Y=treatment means.
<sup>2</sup> Calibration Curve-Basal Diet phosphorus: Calculated by setting the regression equations for P (treatments 1 to 3) equal with those created for phytase followed by subtracting phosphorus content of basal diet
<sup>3</sup>Positive Control-Negative Control: Calculated by comparing phosphorus contents of positive and negative controls.

Regardless of the method of calculation and different response criteria, there was significant difference between strains (P<0.05). Phosphorus equivalences calculated for layer hens were significantly higher than broiler chickens. Similar observations have been reported by van der Klis et al., (1997), who reported a greater phytase efficacy in layer hens compared to those were reported by Camden et al., (2001) and Tamim et al., (2004) in broiler chickens. Leske and Coon (1991) have stated that, this might be due to the longer retention time of digesta in gastrointestinal tract of layers than broilers. On the other hand, the extent of phytase activity is not only a function of retention time of digesta in forestomach tract, but also the phosphorus content of basal diet can influence the response of bird to the phytase. In current study, the lower AvP content of basal diet and the nature of basal resulted in higher P equivalence of phytase in layer hens than broilers.

Zaghari (2009) showed that formulating of diet using the claimed nutrient equivalence of a commercial enzyme resulted in different responses in broiler chickens compared to layer hens. Totally, results of current study have shown that there are some interfering factors such as strain of bird, considering basal diet AvP or ignoring it, and method of calculation, which result in significant differences between evaluated P equivalence of a specific phytase. Recommendation of a single P equivalence for all strains and diet types is ambiguous for the end user to include the matrix value of enzyme claimed by the supplier in diet formulation. Despite of the AvP content of basal diet which resulted in differences between layer hens and broilers, method of estimating of P equivalence of phytase seems to be determinant in calculating P equivalence of phytase in any given strain.

## CONCLUSIONS ANDAPPLICATIONS

1. Results of this experiment demonstrated that the average P equivalence value of phytase (300-500 FTU/kg) for BWG ranged from 0.186 to 0.385 in CC method, 0.094 to 0.170 in CC-BD method and 0.100 to 0.106 in CNP method for broiler chickens.
2. In layer hens the lowest value obtained for 500 FTU/kg phytase was seen in CNP method (0.304) and CC method calculated the highest equivalency value (0.564).
3. The method of calculation which subtracted basal diet P content form total P released by phytase, yielded the more reliable P equivalence than CC. Adopting this method of calculation and BWG as a response criteria in CC-BD method, phytase levels of 300, 400 and 500U/kg of diet were equivalent to the addition of 0.100, 0.131 and 0.161%P from mono-calcium phosphate in 14- to 42-d-old broilers.
4. There was significant difference between different methods and even between two subclasses of a major method of calculation (i.e. calibration curves of performance response) of P equivalence of phytase.
5. There was significant difference between various strains (broilers and layers) in terms of P equivalence values.
6. Different traits had no significant influence on P equivalence of phytase.

## Ethics statement

All procedures including animal welfare, husbandry and experimental procedures were evaluated and approved by the Institutional Animal Care and Ethics Committee of the Iranian Council of Animal Care (Care ICoA 1995).

## Data availability statement

The data that support the findings of this study are available from the corresponding author, upon reasonable request, subject to restrictions and conditions.

## REFERENCES

Abd El-Hack, M. E., Alagawany, M., Arif, M., Emam, M., Saeed, M., Arain, M. A., Siyal, F. A., Patra, A., Elnesr, S. S., and Khan, R. U. 2018. The uses of microbial phytase as a feed additive in poultry nutrition – a review. Ann. Anim. Sci. 18:639–658.

Adedokun, S. A., Sands, J. S., and Adeola, O. 2004. Determining the equivalent phosphorus released by an Escherichia coli-derived phytase in broiler chicks. Can. J. Anim. Sci. 84:437–444.

Bedford, M. R., and Cowieson, A. J. 2020. Matrix values for exogenous enzymes and their application in the real world. J. Appl. Poult. Res. 29:15–22.

Bedford, M. R., and Patridge, G.G. 2011. Factors influencing phytase efficacy. Pages 187-190 in Enzymes in farm animal nutrition. 2nd ed, CAB International, London, UK.

Bedford, M. R., Walk, C. L., Masey O’Neill, H. V. 2016. Assessing measurements in feed enzyme research: Phytase evaluations in broilers. J. Appl. Poult. Res. 25: 305–314.

Camden, B. J., Morel, P. C. H., Thomas, D. V., Ravindran, V., and Bedford, M. R. 2001. Effectiveness of exogenous microbial phytase in improving the bioavailabilities of phosphorus and other nutrients in maize-soybean diets for broilers. Anim. Sci. 73:289–297.

Denbow, D. M., Ravindran, V ., Kornegay, E.T. Yi, T., and Hulet, R. M. 1995. Improving phosphorus availability in soybean meal for broilers by supplemental phytase. Poult. Sci. 74:1831–1842.

Dersjant-Li, Y., Hruby, M., Evans, C., and Greiner, R. 2019. A critical review of methods used to determine phosphorus and digestible amino acid matrices when using phytase in poultry and pig diets. J. Appl. Anim. Nutr.7:1–9.

Farrell, D. J., Martin, E., Preez, J. J., Bongarts, M., Sudaman, A., and Thomson, E. 1993.The Beneficial Effects of a Microbial Phytase in Diets of Broiler Chickens and Ducklings.J. Anim. Physiol. An. N. 69:278–283.

Fernandez, S. R., Charraga, S., and Avila-Gonzalez, E. 2019. Evaluation of a new generation phytase on phytate phosphorus release for egg production and tibia strength in hens fed a corn-soybean meal diet. Poult. Sci. 98:2087–2093.

Francesch, M., Broz, J., and Brufau, J. 2005. Effects of experimental phytase on performance, egg quality, tibia ash content and phosphorus bioavailability in laying hens fed on maize-or barley-based diets. Brit. Poult. Sci. 46:340–348.

Ghosh, A., Mandal, G. P., Roy, A., and Patra, A. K. 2016. Effects of supplementation of manganese with or without phytase on growth performance, carcass traits, muscle and tibia composition, and immunity in broiler chickens. Livest. Sci. 191:80–85.

Han, J. C., Yang, X. D., Qu, H. X., Xu, M., Zhang, T., Li, W. L., Yao, J. H., Liu, Y. R., Shi, B. J., Zhou, Z. F. and Feng, X. Y. 2009. Evaluation of equivalency values of microbial phytase to inorganic phosphorus in 22-to 42-day-old broilers. J. Appl. Poult. Res. 18:707–715.

Igbasan, F. A., Männer, K., Miksch, G., Borriss, R., Farouk, A., and Simon, O. 2000. Comparative studies on the in vitro properties of phytases from various microbial origins. Arch. Anim. Nutr. 54:353–373.

Jalal, M. A., and Scheideler, S. E. 2001. Effect of supplementation of two different sources of phytase on egg production parameters in laying hens and nutrient digestibility. Poult. Sci. 80:1463–1471.

Jendza, J. A., Dilger, R. N., Sands, J. S., and Adeola, O. 2006. Efficacy and equivalency of an Escherichia coli-derived phytase for replacing inorganic phosphorus in the diets of broiler chickens and young pigs. J. Anim. Sci. 84:3364–3374.

Keshavarz, K. 2000. The effect of different levels of nonphytate phosphorus with and without phytase on the performance of four strains of laying hens. Poult. Sci. 82:71–91.

Leske, K., and Coon, C. N. 1999.A bioassay to determine the effect of phytase on phytate phosphorus hydrolysis and total phosphorus retention of feed ingredients as determined with broilers and laying hens. Poult. Sci. 78:1151–1157.

Li, W., Angel, R., Kim S. W., Jimenez-Moreno, E., Prosz kowiec-Weglarz, M., and Plumstead, P. W. 2015. Age and adaptation to Ca and P deficiencies: 2. Impacts on amino acid digestibility and phytase efficacy in broilers. Poult. Sci. 94:2917–2931.

Li, W., Angel, R., Proszkowiec-Weglarz, M., Kim, S. W., Jiménez-Moreno, E., and Plumstead, P. W. 2013. Criteria of response and Ca concentration affect estimates of phytase equivalence to monocalcium phosphate. Poult. Sci. 92:48.

Liu, N., Ru, Y. J., Li, F. D., Wang, J., and Lei, X. 2009. Effect of dietary phytate and phytase on proteolytic digestion and growth regulation of broilers. Arch. Anim. Nutr. 63:292–303.

Nahm, K. H. 2002. Efficient feed nutrient utilization to reduce pollutants in poultry and swine manure. Environ. Sci. Technol. 32:1–16.

Näsi, M., Partanen, K. and Piironen, J. 1999. Comparison of Aspergillusniger phytase and Trichodermareesei phytase and acid phosphatase on phytate phosphorus availability in pigs fed on maize-soybean meal or barley-soybean meal diets. Arch. Anim. Nutr. 52:15–27.

Newkirk, R. W., and Classen, H. L. 2001.The non-mineral impact of phytate in canola meal fed to broiler chicks. Anim. Feed Sci. Technol. 9:115–128.

Plumstead, P., Sriperm, N., and Swann, D. 2013. Modeling effects of Buttiauxella phytase on energy and amino acid utilisation in broilers. International Poultry science Forum, Atlanta, USA.

Potter, L. M. 1988. Bioavailability of phosphorus from various phosphates based on body weight and toe ash measurements. Poult. Sci. 67:96–102.

Qian, H., Kornegay, E. T., and Denbow, D. W. 1997. Utilization of phytate phosphorus and calcium as influenced by microbial phytase, cholecalciferol, and the calcium:total phosphorus ratio in broiler diets. Poult. Sci. 76:37–46.

Ravindran, V., Cabahug, S., Ravindran, G., and Bryden, W. L. 1999. Influence of microbial phytase on apparent ileal amino acid digestibility of feedstuffs for broilers. Poult. Sci. 78:699–706.

Ravindran, V., Cabahug, S., Ravindran, G., Selle, P. H., and Bryden, W. 2000. Response of broiler chickens to microbial phytase supplementation as influenced by dietary phytic acid and non-phytate phosphorous levels. II. Effects on apparent metabolisable energy, nutrient digestibility and nutrient retention. Brit. Poult. Sci. 41:193–200.

Ribeiro, Jr. V., Salguero, S. C., Gomes, G., Barros, V. R. S. M., Silva, D. L., Barreto, S. L. T., Rostagno, H. S., Hannas, M. I., and Albino, L. F. T. 2016. Efficacy and phosphorus equivalency values of two bacterial phytases (Escherichia coli and Citrobacterbraakii) allow the partial reduction of dicalcium phosphate added to the diets of broiler chickens from 1 to 21 days of age. Anim. Feed. Sci.Technol. 221:226–233.

Rodriguez, E., Han, Y., and Lei X. G. 1999b. Different sensitivity of recombinant Aspergillusniger phytase (r-PhyA) and Escherichia coli pH 2.5 acid phosphatase (r-AppA) to trypsin and pepsin in vitro. Arch. Biochem. .365:262–267.

Rodriguez, E., Han, Y., and Lei, X. G. 1999a. Cloning, sequencing, and expression of an Escherichia coli acid phosphatase/phytase gene (appA2) isolated from pig colon. Biochem. Bioph. Res. Co. 257:117–123.

Rutherfurd, S. M., Chung, T. K., and Moughan, P. J. 2004.The effect of a commercial microbial phytase preparation on the in vitro release of phosphorus and amino acids from selected plant feedstuffs supplemented with free amino acids. J. Anim. Feed Sci. 13:677–690.

Rutherfurd, S. M., Chung, T. K., Morel, P. C. H., and Moughan, P. J. 2004. Effect of microbial phytase on the ileal digestibility of phytate phosphorus, total phosphorus, and amino acids in a low phosphorus diet for broilers. Poult. Sci. 83:61–68.

Selle, P., Ravindran, V. 2007. Microbial phytase in poultry nutrition. Anim. Feed Sci. Technol. 135:1–41.

Shet, D., Ghosh, J., Ajith, S., and Awachat, V. B. 2017. Efficacy of dietary phytase supplementation on laying performance and expression of osteopontin and calbindin genes in eggshell gland. Anim.Nutr.4: 52–58.

Simons, P. C. M., and Versteegh, H. A. J. 1992. Informative study concerning the effect of the addition of microbial phytase to layer feed. Spelderholt, publication no. 573 (NL).

Simons, P. C. M., and Versteegh, H. A. J. 1993. Role of phytase in poultry nutrition. Pages 181–186 in: C. Wenk, and M. Boessinger, ed. Proc.1st Symposium of Enzymes in Animal Nutrition. Kartause, Ittingen, Switzerland.

Tamim, N. M., Angel, R., and Christman, M. 2004. Influence of dietary calcium and phytase on phytate phosphorus hydrolysis in broiler chickens. Poult. Sci. 83:1358–67.

Tran, T. T., Hatti-Kaul, R., Dalsgaard, S., and Yu, S. 2011. A simple and fast kinetic assay for phytases using phytic acid–protein complex as substrate. Anal. Biochem. 410:177–184.

Um, J. S., and Paik, I. K. 1999. Effects of microbial phytase supplementation on egg production, eggshell quality, and mineral retention of laying hens fed different levels of phosphorus. Poult. Sci. 78:75–79.

Van der Klis, J. D., Versteegh, H. A., Simons, P. C., and Kies, A. K. 1997. The efficacy of phytase in corn-soybean meal-based diets for laying hens. Poult. Sci. 76:1535–1542.

Vieira, S. L., Anschau, D. L., Stefanello Serafini, N.C., Kindlein, L., Cowieson, A. J., and Sorbara, J. O. B. 2015. Phosphorus equivalency of a Citrobracterbraakii phytase in broilers. J. Appl. Poult. Res. 00:1–8.

Waldroup, P. W. 1999. Nutritional approaches to reducing phosphorus excretion by poultry. Poult. Sci. 78:683–691.

Whitehead, C. C., and Fleming, R. H. 2000. Osteoporosis in cage layers. Poult. Sci. 79:1033–1041.

Wu, Y. B., Ravindran, V., Morel, P. C. H., Hendriks, W. H., and Pierce, J. 2004. Evaluation of a microbial phytase, produced by solid-state fermentation, in broiler diets. 1. Influence on performance, toe ash contents, and phosphorus equivalency estimates. J. Appl. Poult. Res. 13:373–383.

Zaghari, M. 2009. Evaluation of using phytase nutrient equivalency values for layer hens and broiler chickens. J. Agr. Sci. and Technol.11:57–66.

Żyła, K. 1993. The role of acid phosphatase activity during enzymic dephosphorylation of phytates by Aspergillus niger phytase. World J. Microb. Biot. 9:117–119.

